# Uneven declines between corals and cryptobenthic fish symbionts from multiple disturbances

**DOI:** 10.1101/2021.01.13.426488

**Authors:** Catheline Y.M. Froehlich, O. Selma Klanten, Martin L. Hing, Mark Dowton, Marian Y.L. Wong

## Abstract

With the onset and increasing frequency of multiple disturbances, the recovery potential of critical ecosystem-building species and their mutual symbionts is threatened. Similar effects to both hosts and their symbionts following disturbances have been assumed. However, we report unequal declines between hosts and symbionts throughout multiple climate-driven disturbances in reef-building *Acropora* corals and cryptobenthic coral-dwelling *Gobiodon* gobies. Communities were surveyed before and after consecutive cyclones (2014, 2015) and heatwaves (2016, 2017). After cyclones, coral size and goby group size decreased similarly by 28-30%. After heatwave-induced bleaching, coral size decreased substantially (47%) and the few gobies recorded mostly inhabited corals singly. Despite several coral species still occurring after bleaching, all goby species declined, leaving 78% of corals uninhabited. These findings suggest that gobies, which are important mutual symbionts for corals, are unable to cope with consecutive disturbances. This disproportionate decline could lead to ecosystem-level disruptions through loss of key symbiotic services to corals.

## INTRODUCTION

Multiple disturbances over short periods can disrupt important processes and threaten the persistence of ecosystems(1, 2). From species survival to population bottlenecks and trophic disruptions, such consecutive disturbances may transform entire environments(1–4). The ability for ecosystems to recover depends on the frequency and intensity of multiple events, which are predicted to increase with climate change(1, 5). Relationships within complex environments can deteriorate in an accelerated fashion as a result(1). Whether organisms persist in the short-term during extreme consecutive disturbances will determine their recovery potential and those of associated organisms(6–8). We need to understand whether ecological relationships are resilient to consecutive disturbances in order to better align future strategies for ecosystem conservation(6, 9).

One such ecological relationship that may prove fragile from consecutive disturbances is mutualism, which occurs in many taxa(6, 9). Mutual symbioses are observed in all environments and even promote life in otherwise inhospitable areas(9). A small shift in environmental conditions may change the nature of such relationships, like mutualism becoming parasitism, or relationships ceasing if one symbiont becomes locally threatened(6). Climate-driven disturbances can lead to breakdowns of mutualisms like those responsible for preventing seagrass degradation(10), maintaining myrmecophyte-dominated savannahs(11), sustaining coral survival(12), and promoting microbe-assisted biodiversity(9, 13). Collapse of mutual symbioses may have flow-on effects by destabilizing habitats and causing deleterious ecosystem consequences(6, 13). Studies need to assess the consequences of disturbances on mutual symbioses in order to predict flow-on effects to ecosystems.

Mutually beneficial taxa are especially vulnerable to climate-driven disturbances when ecosystem building blocks are threatened, as seen in coral reefs(2). As the frequency and intensity of storms and heatwaves are increasing, corals are being exposed to disturbances in rapid succession(14–16). Up to 11% of coral reef fishes depend on live corals for survival through food, settlement and shelter(17, 18). In return, coral-associated fishes promote coral resilience by reducing disease, algal growth, and increasing nutrient cycling(19–23). However, disturbance studies are largely focused on corals(14–16). If fish symbionts decline disproportionately from climate-driven impacts(24), then corals will be exposed to additional threats as there is little functional overlap in coral reefs(25). Disproportional declines in corals and their mutualistic symbionts may lead to ecosystem shifts(26) if consecutive disruptions become the norm(5).

Here, we examined the impacts of multiple climate-driven disturbances on the persistence of coral-fish symbioses using the most susceptible reef-building corals (genus *Acropora*)(16, 27) and their mutually beneficial inhabitants, cryptobenthic coral-dwelling gobies (genus *Gobiodon*)(20, 21). In return for shelter, breeding sites and food from corals(28, 29), gobies remove harmful seaweed, deter corallivores, and increase nutrient cycling(19–21) (Fig 1a). Gobies are often overlooked in disturbance studies, even though as cryptobenthic fishes they are critical to the trophic structure of coral reefs(4). We surveyed coral and goby communities throughout four consecutive disturbances at Lizard Island, Great Barrier Reef, Australia. Within four years, the reef experienced two cyclones (2014, 2015), and two unprecedented heatwaves that caused widespread bleaching (2016, 2017)(30). Our study quantified the additive impacts of cyclones and heatwaves on the persistence of corals and their goby symbionts over a short space of ecological time.

**Fig 1.**
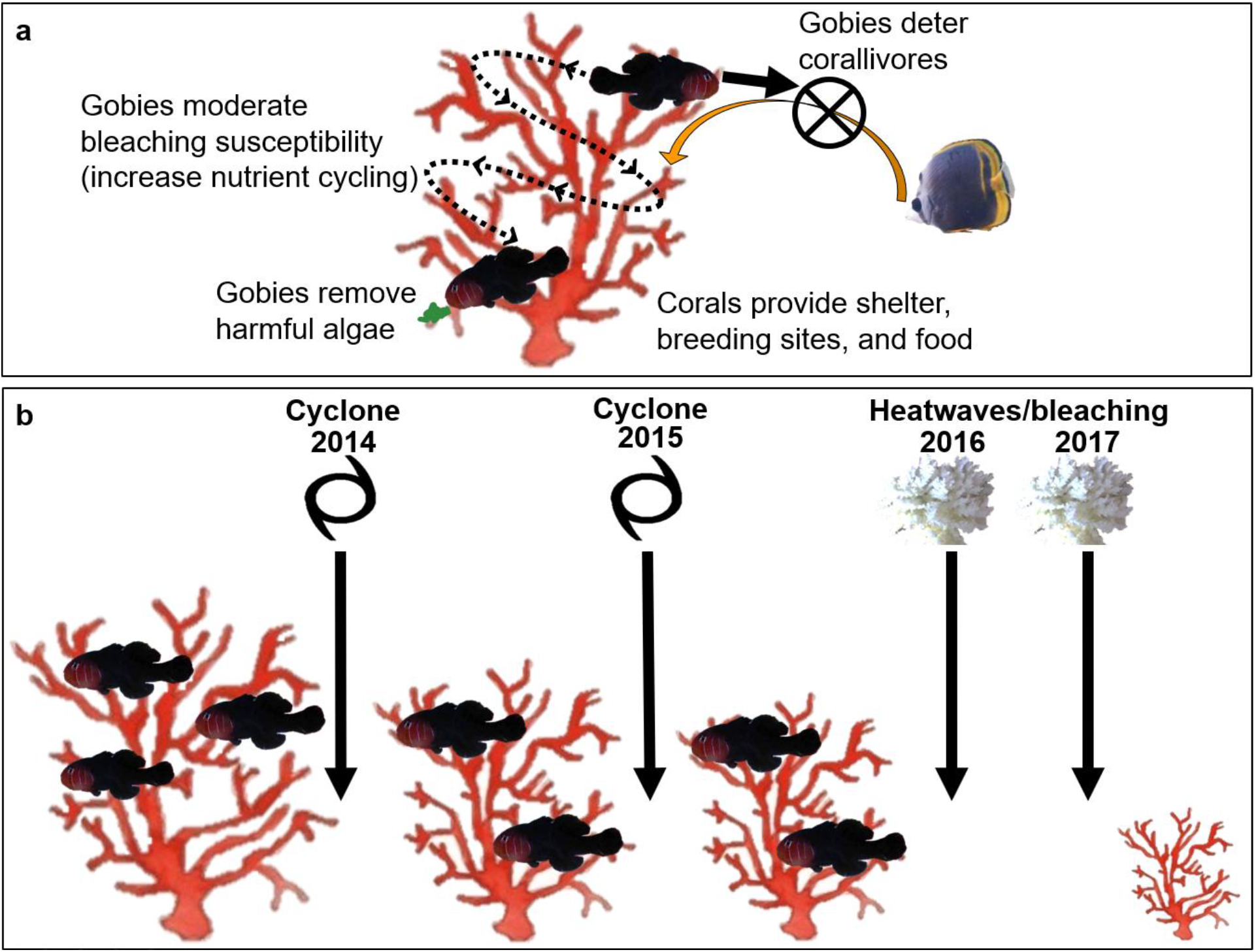
Drastic shifts to the mutual symbiosis of corals and cryptobenthic coral-dwelling gobies following multiple disturbances. **A** Benefits that each symbiont receives from the mutual symbiosis(19–21, 28, 29). **B** Summary of the findings highlighting changes to corals and gobies from each consecutive disturbance with coral-dwelling gobies from the genus *Gobiodon* in scleractinian corals from the genus *Acropora*. Reductions in coral size are drawn to scale and relative to changes in means among disturbances.

## RESULTS AND DISCUSSION

### Host and mutual symbionts decline at different rates following consecutive cyclones and bleaching

In February 2014, prior to cyclones and bleaching events, most *Acropora* corals were inhabited by *Gobiodon* coral gobies. Gobies were not found in corals under 7-cm average diameter, therefore we only sampled bigger corals. The vast majority of transects (95%) had *Acropora* corals. On average there were 3.24 ± 0.25 (mean ± standard error) *Acropora* coral species per transect (Fig 2a) and a total of 17 species. Average coral diameter was 25.4 ± 1.0 cm (Fig 2b), with some corals reaching over 100 cm. Only 4.1 ± 1.4 % of corals lacked any goby inhabitants (Fig 2c). On average there were 3.37 ± 0.26 species of gobies per transect (Fig 2d) and a total of 13 species. In each occupied coral there were 2.20 ± 0.14 gobies (Fig 2e), with a maximum of 11 individuals of the same species.

**Fig 2.**
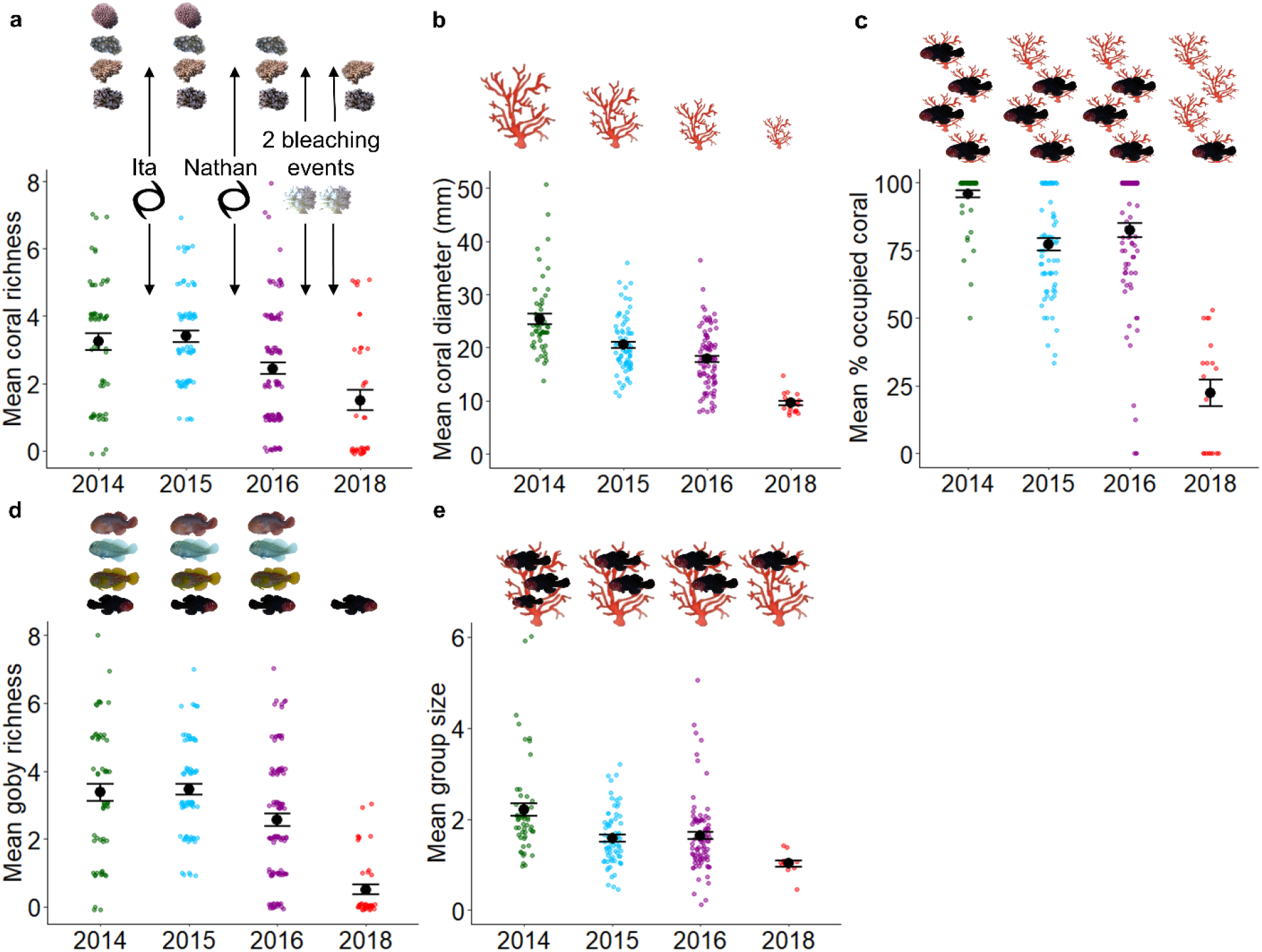
Effects of consecutive climate disturbances on coral and goby populations. Changes in *Acropora* **a** richness (n = 279), and **b** average diameter (n = 244), **c** percent goby occupancy (n = 244) and *Gobiodon* **d** richness (n=279), and **e** group size (n = 230) before and after each cyclone (black cyclone symbols) and after two consecutive heatwaves/bleaching events (white coral symbols) around Lizard Island, Great Barrier Reef, Australia. Error bars are standard error. Fish and coral symbols above each graph illustrate the change in means for each variable among sampling events from post-hoc tests.

In January-February 2015, 9 months after Cyclone Ita (category 4) struck from the north (Supplementary Fig 1), follow-up surveys revealed no changes to coral richness (p = 0.986, see Supplementary Table 1 for all statistical outputs) relative to February 2014, but corals were 19% smaller (p < 0.001, Fig 2a-b). Cyclonic activity may have damaged existing corals(31), which might explain smaller corals. Alternatively, corals may have died from cyclonic damage(31), but previously undetected corals (less than 7-cm average diameter threshold for surveys) may have grown and accounted for finding smaller corals and no changes to species richness. After the cyclone, gobies occupied 76% of live corals, which meant that occupancy dropped by 19% (p < 0.001, Fig 2c). Goby richness did not change after the first cyclone relative to February 2014 (p = 0.997, Fig 2d). However, goby group sizes were 28% smaller (p < 0.001), with gobies mostly occurring in pairs, and less so in groups (Fig 2e). Smaller groups were likely due to their coral hosts being smaller than before the cyclones as there is an indirect link between group size and coral size(32).

In January-February 2016, 10 months after Cyclone Nathan (category 4) struck from the south (Supplementary Fig 1), our follow-up surveys revealed 26% fewer coral species (p = 0.008), and 13% smaller corals (p = 0.029) relative to February 2015 (Fig 2a-b). Many corals were damaged, and bigger corals were likely heavily damaged and reduced in size. As *Acropora* corals vary in several morphological traits such as branch thickness, such characteristics might alter their susceptibility to cyclonic damage(31, 33) and likely explain a decrease in coral richness. There was no change to coral occupancy relative to February 2015 (p = 0.167, Fig 2c). Goby richness however did not mirror declines to their coral hosts as there was no change relative to February 2015 (p = 0.060, Fig 2d). Goby group size did not change relative to February 2015 and most individuals occurred only in pairs (p = 1.000, Fig 2e). Since the second cyclone did not add additional changes to coral occupancy, goby richness or goby group size, gobies may have exhibited some ecological memory(30) from the first cyclone. However, when combining the effects of consecutive cyclones, coral and goby symbioses were disrupted substantially. Coral hosts were 30% smaller relative to 2014 (pre-disturbances), 25% of hosts were uninhabited compared to only 4% in 2014, and gobies were no longer living in groups, instead living in pairs (Fig 1b). These acute disturbances had effects lasting longer than 10 months and will likely require many years to return to pre-disturbance status(14).

Unfortunately, there was no time for recovery from cyclones before two prolonged heatwaves caused widespread bleaching in March-April 2016 and February-May 2017 (Supplementary Fig 1). Ten months after the second bleaching event (Jan-Feb 2018), we returned to Lizard Island and rarely found live corals along our transects. Half (50%) of the transects lacked any living *Acropora* corals compared to just 5% before any disturbance (2014). There were 39% fewer coral species (p = 0.009) relative to February 2016, with only 1.5 ± 0.31 species per transect (Fig 2a). Corals were 47% smaller than in February 2016 (p < 0.001, Fig 1b&2b), averaging 9.57 ± 0.39 cm coral diameter (maximum 21 cm). Acroporids were also the most susceptible family to bleaching from these back-to-back heatwaves across the Great Barrier Reef and their coral recruitment was at an all-time low(2, 16). Since corals were lethally bleached during the prolonged heat stress only a few acroporids species survived these consecutive events(34). The 2015-2016 heatwave caused extensive bleaching in many areas around the world, which has become a globally common phenomenon in the last few decades(5, 35).

After consecutive heatwaves, coral gobies faced even more drastic declines than their coral hosts in all our survey variables. Of the few live corals recorded, most (77.7 ± 4.8%) corals lacked gobies compared to just 4% pre-disturbance (2014), and 24% after cyclones (p < 0.001, Fig 2c). For the first time, only after heatwaves, we observed a change in goby richness with 80% fewer goby species per transects relative to February 2016 (p < 0.001, Fig 2d), even though consecutive cyclones did not affect goby richness. Alarmingly, gobies were no longer found in groups (p = 0.036), rarely in pairs (n = 3), and the few observed occurred singly (Fig 2e). For these long-living and nest brooding fishes(28, 36), finding gobies predominantly without mates suggests that reproduction likely ceased or was significantly delayed for most individuals in the population(28). An interruption in mate pairing likely led to extremely low recruitment and turnover rates in gobies from climatic disturbances.

Gobies declined substantially more than coral hosts after consecutive heatwaves, leaving most corals uninhabited (Fig 1b). Although communities still had not recovered from cyclonic disturbances before prolonged heatwaves, we suspect that heatwaves had more devastating impacts on gobies than cyclones. Gobies have a strong tendency to stay in the same coral they settle in as recruits(37) as long as the coral is alive(38), yet many may have unsuccessfully attempted to find other corals once their coral was lethally bleached(4). Unlike gobies, other coral-dwelling fishes, like damselfish recruits, successfully adopted alternative habitat, including dead corals(39). Gobies did not adopt alternative habitat and were surprisingly absent from most living corals.

Importantly, goby richness did not change after consecutive cyclones and only changed after heatwaves. Thus coral host death likely is not the only stressor and gobies may have suffered physiological consequences from prolonged heatwaves(40–42). Although gobies can survive short exposures of hypoxia(43), extended periods of reduced wind-induced mixing and thermal stress may jeopardize physiological functioning(44, 45). Indeed, reef fishes can lose the ability to detect predators, kin, and habitat(40–42), and to reproduce from thermal stress(45, 46). Gobies likely lost similar functioning from heatwaves leading to high mortality and little goby turnover, which left many healthy corals unoccupied. A lack of mutual goby symbionts following consecutive disturbances suggests that coral hosts may begin exhibiting additional threats to their recovery(19–21). Such declines and potential physiological consequences may also hold true for other coral-dwelling organisms, like symbiotic xanthid crabs(47). Since acroporid corals are important building blocks for coral reef ecosystems, greater declines in their symbionts from multiple disturbances may reduce the persistence of corals and destabilize habitats over large scales.

### Communities of goby symbionts exhibit greater changes than communities of coral hosts from multiple disturbances

In February 2014, before the consecutive climatic events, we recorded 17 species of *Acropora* corals, with the most common being *A. gemmifera, A. valida, A. millepora, A. loripes, A. nasuta, A. intermedia, A. tenuis,* and *A. cerealis*. Thirteen species of *Gobiodon* gobies were recorded, with the most common being *G. rivulatus, G. fuscoruber, G. brochus, G. histrio, G. quinquestrigatus,* and *G. erythrospilus*. Each disturbance changed the assemblages of both corals (p < 0.001, Fig 3a) and gobies (p < 0.001, Fig 3b), yet the changes in both corals and gobies did not mirror each other (Fig 3).

**Fig 3.**
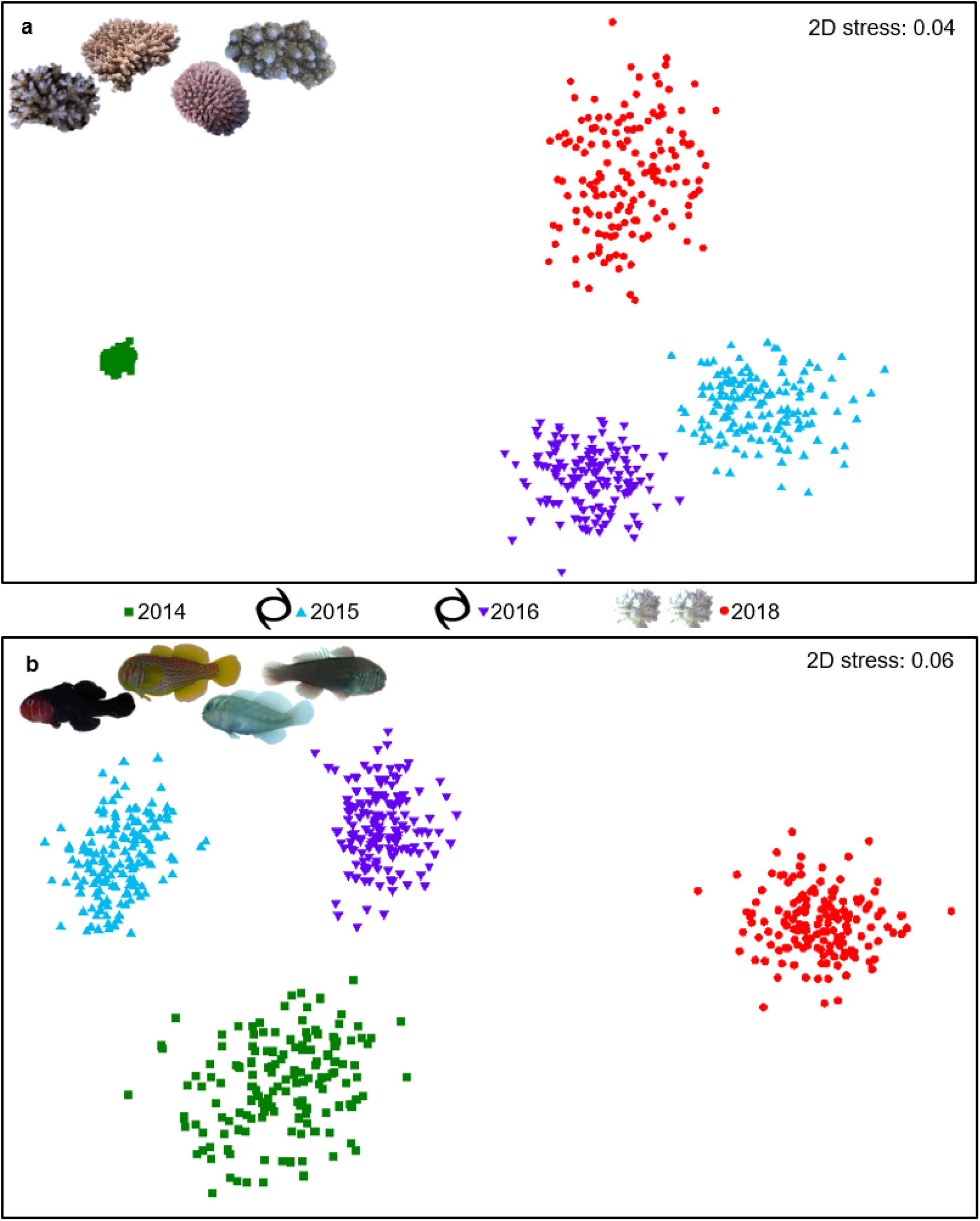
Shifts in communities of corals and gobies throughout consecutive climate disturbances. The changes in communities before and after each cyclone (black cyclone symbols) and after two consecutive heatwaves/bleaching events (white coral symbols) at Lizard Island, Great Barrier Reef, Australia, for **a** *Acropora* corals (n = 279) and **b** *Gobiodon* gobies (n = 279) visualized on non-metric multidimensional scaling plots. Each colored point represents a single transect, black points represent bootstrapped averages (avg), and points closer together are more similar in species composition than points further apart.

After the first cyclone, 11 *Acropora* species were found, and the common species were recorded more often relative to February 2014 (p = 0.009, Fig 3a&4a). Previously rare species, *A. cerealis* and *A. valida* became more often recorded as well. However, *Acropora intermedia*, which was previously recorded in several transects, was no longer observed; this is likely due to its branches being long and thin, thus highly susceptibility to damage(31). Goby assemblages were also altered after the first cyclone (p = 0.003, Fig 3b), and some common species were more often recorded while others made up less of the samples relative to 2014 (Fig 4b). *Gobiodon histrio* and *G. rivulatus* were recorded more often compared to others, and so were their preferred hosts, *A. nasuta* and *A. gemmifera*, respectively (Fig 4)(48). However, *Gobiodon fuscoruber* was recorded less frequently even though its common host, *A. millepora*(48), was recorded more frequently than other corals (Fig 4). Unfortunately, two rare gobies were no longer recorded (*G. citrinus* and *G. okinawae*), and both preferred *A. intermedia*(48), which also disappeared.

**Fig 4.**
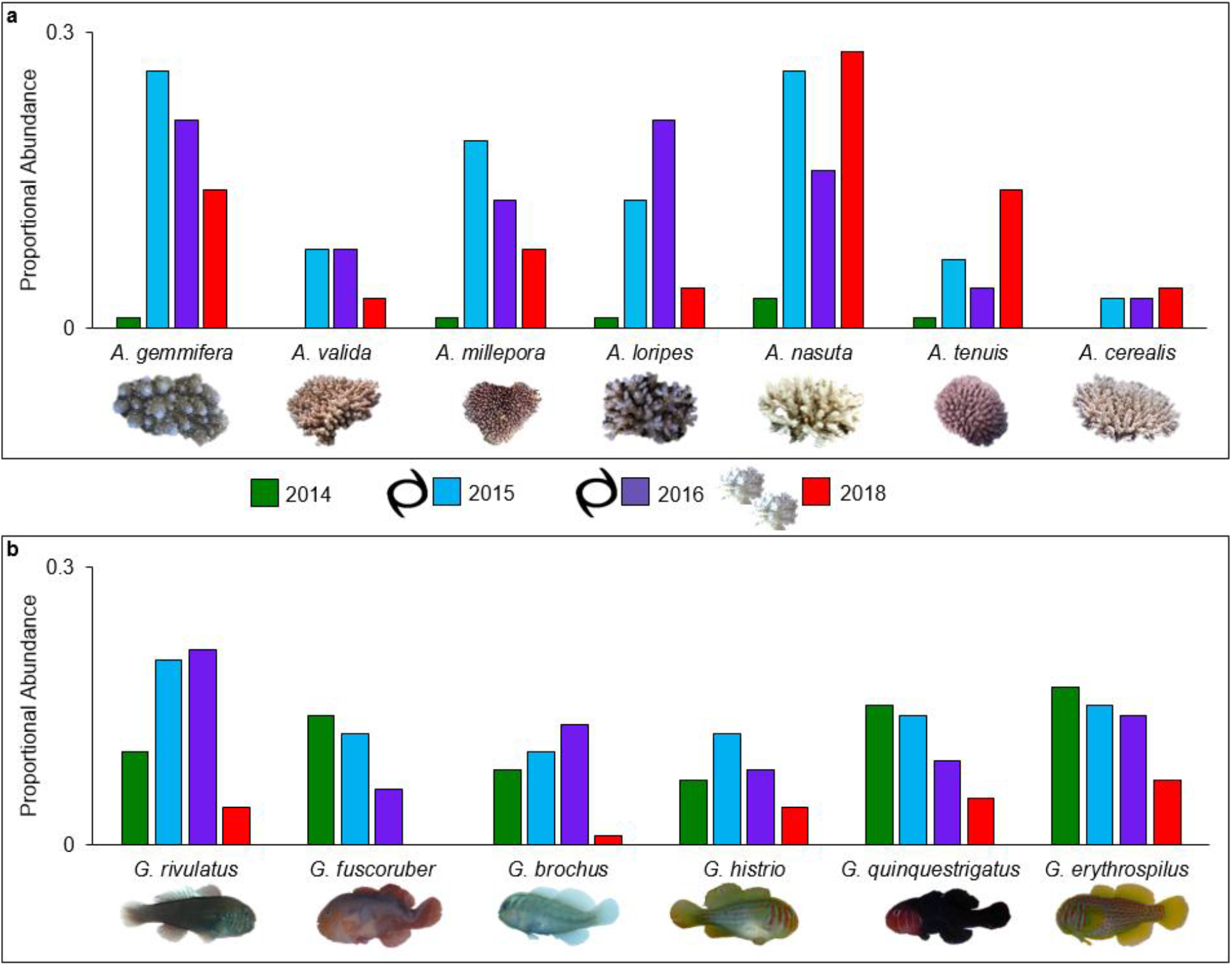
Changes in abundance of coral and goby species before and after each consecutive climate disturbances. The proportional abundances for the most common species within each sample for **a** *Acropora* corals (n = 279) and **b** *Gobiodon* gobies (n = 279) is shown before and after each disturbance around Lizard Island, Great Barrier Reef, Australia: effect of each cyclone (black cyclone symbols) and combined effect of two consecutive heatwaves/bleaching events (white coral symbols). Standardized abundances were calculated from similarity percentage analyses for each sampling event.

After the second cyclone, we found mixed results in coral assemblages (p < 0.001, Fig. 3a). Although 15 *Acropora* species were found after the second cyclone (5 more than after the previous cyclone) and no species were locally extirpated, only *A. loripes* became more common (Fig 4a). Several of the most common corals (i.e. *A. gemmifera, A. millepora, A. nasuta, A. tenuis*) became less reported after the second cyclone (Fig 4a). Goby communities were altered once again (p < 0.001, Fig 3b), this time with fewer species increasing in proportion, and more species decreasing (Fig 4b). However, all *Gobiodon* species were encountered, even *G. citrinus* and *G. okinawae* that originally disappeared after the first cyclone. *Gobiodon brochus* increased in proportion and so did its common host *A. loripes*(48). However, *G. rivulatus* increased even though its preferred host *A. gemmifera* decreased (Fig 4)(48).

After consecutive bleaching events, the reef was left with few, very small corals. Although the coral community after bleaching was distinct from each disturbance time point (p < 0.001), all disturbed communities grouped together compared to the pre-disturbance community (2014, Fig 3a). After bleaching, the most species were recorded (22 in total). A few *A. intermedia* were again recorded after none were observed following the first cyclone, along with 9 rare and previously unrecorded *Acropora* species. However, some species were no longer observed, e.g. *A. 10ivaricate* (previously rare)*, A. granulosa* (previously rare), and *A. humilis* (previously common). Many of the common coral species became rare after bleaching (Fig 4a). In coral reefs, *Acropora* are one of the most susceptible coral genera to cyclone damage and bleaching in a warming climate(16, 31), which explains such steep declines in many *Acropora* species. Surprisingly, *A. cerealis*, which was previously rare, had since become more common in our surveys despite multiple disturbances (Fig 4a). In other areas though, such as the Andaman Bay, *A. cerealis* was one of the most lethally bleached species(35). Regional differences in thermal plasticity and coral recruitment may have disproportionately affected the survival thresholds of identical species.

For coral gobies though, there were even more dramatic effects of consecutive bleaching than for corals. Goby communities were the most distinct after bleaching (p < 0.001), while all previous surveys grouped together (Fig 3b). Every goby species declined after bleaching, and half of the species were no longer recorded (Fig 4b). Some species were locally extirpated, including *G. citrinus* (previously rare), *G. sp. D* (previously rare), *G. bilineatus* (previously common), and *G. fuscoruber* (previously common). None of the locally extirpated species were observed during random searches. Only 6 species remained, and no previously unrecorded species were observed. As expected, gobies were never found in dead corals, as they can only survive in live corals (albeit surviving in stressed corals(38)). These findings highlight the greater impact that multiple disturbances have on symbiont communities, especially when disturbances are a mix of acute (short-term) and prolonged (long-term) events. We observed a loss of biodiversity for gobies from multiple disturbances, whereas their coral hosts were more diverse even though fewer corals were recorded.

The study demonstrates the effects that multiple disturbances have on reef ecosystems down to the level of crucial mutual symbioses. Disturbance studies have primarily focused on the effects to corals(16, 30, 31), yet cryptobenthic fishes are often overlooked(4). We may be missing bigger effects of disturbances on coral reef ecosystems, especially since cryptic fishes make up a large part of reef biodiversity and are crucial prey for many taxa(4). This study is one of few long-term studies to record species-level changes in cryptobenthic fishes from multiple consecutive disturbances. Intriguingly, although corals and gobies responded similarly at first to the initial two cyclones, they then diverged in their responses after additional stress from heatwaves. Here we show that gobies declined faster on a community and species level than their coral hosts, which will likely leave corals exposed to algal growth, poor nutrient cycling, and corallivory(19–21) (Fig 1). The unwillingness of gobies to use alternative habitat in the short-term may drastically reduce their resilience to disturbances, threatening localized extinction(49).

Declines from a single disturbance have the potential for a resilience, but multiple events will require long-term recovery(31, 32) as most corals are uninhabited after consecutive disturbances (Fig 1b). Although the disturbances in this study were compounded, heatwaves may have had an even stronger effect on gobies since goby communities differed the most after the heatwaves, whereas coral communities remained similarly diverse after each disturbance. Without the added benefits of gobies, surviving corals will likely experience further threats to survival(19–21). Multiple disturbances may even cause ecosystem shifts when the building blocks of the environment, such as hard corals, face extreme declines(6). If mutual symbionts show greater declines than corals, important processes may be exacerbated, further jeopardizing the recovery potential of ecosystem building blocks.

### Future implications for symbiotic relationships from multiple disturbances

Our study demonstrates that consecutive disturbances result in uneven declines between mutual symbionts, and surviving hosts may face additional threats if their mutual and cryptic inhabitants disappear. As climate-driven events becomes more frequent(5), the threat of multiple disturbances will likely deteriorate our ecosystems as mutual symbioses break down(1, 6, 13, 24). Weather events are more common and severe(14), corals are bleaching on a global scale(5), and fires and droughts are becoming more frequent and widespread(1). Although the length and type of the disturbance play important roles in disturbance impacts, few studies have examined the effect of multiple disturbances(30, 31, 50). If successive threats become the norm, a system will already be stressed before a second event strikes, leading to greater consequences(31). Population bottlenecks will inevitably follow(3) and threaten the survival of many organisms globally(7). Flow-on effects will affect closely-associated organisms, especially for those that depend on feedback loops with symbionts(6). In each ecosystem, species are responding differently to disturbances, and mutually beneficial relationships are being tested(6). Our study suggests that multiple disturbances will likely leave ecosystem builders exposed to additional threats if their cryptic symbionts fail to recover.

## METHODS

### Study Location and Sampling Effort

The study was completed at Lizard Island, Great Barrier Reef, Queensland, Australia (14° 40.729’ S, 145° 26.907’ E, Supplementary Fig 1). Four climatic events affected Lizard Island from 2014 to 2018. Cyclone Ita hit in April 2014, and Cyclone Nathan hit in March 2015 (Supplementary Fig 1). The following year 2016, the first extensive mass-bleaching event spanned March to April, and a second extensive mass-bleaching event spanned February to May 2017. A total of 17 sites were first visited in February 2014 before climatic events. After the first cyclone 10 sites were revisited in January-February 2015, 15 sites in January-February 2016 (after second cyclone), and 17 sites in February-March 2018 (after back-to-back heatwaves).

### Survey Method

At each site, goby and coral communities were surveyed within 1 m on either side of 30-m line transects by scuba divers in 2014 (n = 59 transects) and 2018 (n = 40). Transects were completed in 2015 (n = 73) and 2016 (n = 107) using cross-transects—two 4-m x 1-m belt transects laid in a cross around a focal colony. In 2018, random searching for up to one hour (in addition to the transects) was also completed in several areas (n = 28 searches) to determine whether goby species that were missing were simply absent from transects or were instead likely locally extirpated from Lizard Island. For all methods, when a live *Acropora* coral was encountered, the coral was identified to species and measured along three dimensions: width, length, and height(29). A bright torch light (Bigblue AL1200NP) was shone in the coral to quantify the number of goby residents and the *Gobiodon* species inhabiting the coral. Gobies were delineated either as adults or recruits depending on their coloration and size.

### Data Analysis

Univariate analyses were completed to assess changes in the following variables throughout disturbances: adult goby species richness, adult goby group size per coral, percent occupied coral, coral species richness, coral diameter (the three coral dimensional measurements were averaged to calculate an average diameter)(29). Abundance measures were not analyzed because of the differences in survey methodology. Goby and coral richness were count data with several zero data points after multiple disturbances. As such richness variables were each analyzed using two-tailed zero-inflated generalized linear mixed model designs (GLMER: using poisson family) amongst the sampling year (fixed factor) and site (random factor). The following variables were continuous variables and as such were analyzed using two-tailed linear mixed model designs (LMER) amongst the sampling year (fixed) and site (random): coral diameter, goby group size, and percent occupied corals. Variables analyzed with LMER were transformed as required to meet normality and homoscedasticity, which were determined using Q-Q plots, histograms, and residuals over fitted plots. Tukey’s tests were used for differentiating between statistically significant levels within factors. For each univariate analysis, outliers were removed if their standard residuals fell outside of 2.5 standard deviation from 0. A maximum of 7 outliers were removed for any given analysis. All analyses were completed in R version 3.5.2 with the following packages: tidyverse, lme4, lmerTest, LMERConvenienceFunctions, piecewiseSEM, glmmTMB, emmeans, DHARMa, and performance.

Community composition was analyzed separately for corals and gobies. To take into account the different survey techniques, samples were standardized to create proportional abundance as follows: for each survey, we divided each count per species by the total abundance of all species. Only adult gobies were included in the analyses. Communities were analyzed with two-tailed permutational analyses of variance (PERMANOVA). Communities were compared against sampling year (fixed factor) and were controlled for site (random factor) with permutational analyses of variance in PRIMER-E software (v7). Type I error was included because of the unbalanced design with uneven transects per year. Community differences were bootstrapped to a 95% region for a total of 150 bootstraps per year and were visualized on non-metric multidimensional scaling plot. When statistical differences were observed, similarity percentage analyses (SIMPER) were performed to determine what species contributed to the differences observed. Species contributions were cut off to the top 75% of species that contributed the most to differences observed. See supplemental table 1 for all statistical outputs of univariate and multivariate analyses.

## ACKNOWLEDGEMENTS

We acknowledge and pay respect to the Dingaal Aboriginal Traditional Owners of Lizard Island, the sacred site upon which we completed our field work. We greatly thank the assistance of the Lizard Island Research Station staff, and especially Anne Hoggett and Lyle Vail. We thank Kylie Brown, Karen Hing, Grant Cameron, Siobhan Heatwole, and Anna Scott for help in data collection. We thank the Australian Coral Reef Society for organizing and funding a Writing Retreat that provided substantial assistance to manuscript writing. We appreciate Daniel “Al” Alder’s help with PRIMER-E figure production. A special thank you to Cristian Rojas and Jennifer Donelson for helpful feedback on the manuscript. The data collection was funded by the Hermon Slade Foundation to MW. The manuscript writing was supported by the Australian Coral Reef Society Writing Retreat Travel Award and University Postgraduate Award from the University of Wollongong to CF. The study was completed under the University of Wollongong Animal Ethics protocol AE1404 and AE 1725 and under research permits issued by the Great Barrier Reef Marine Park Authority (G13/36197.1 and G15/37533.1).

## DATA AVAILABILITY

Datasets are available at the Knowledge Network for Biocomplexity repository with identifier: *doi:10.5063/5T3HW8*

## AUTHOR CONTRIBUTIONS

C.F. – conceptual framework, data collection, analysis, manuscript writing

O.K. – conceptual framework, data collection, manuscript editing

M.H. – data collection, manuscript editing

M.D. – manuscript editing

M.W. – conceptual framework, data collection, manuscript editing

