## Supplemental Figure and Table for "Uneven declines between corals and cryptobenthic fish symbionts from multiple disturbances"

**SUPPLEMENTARY MATERIAL**
**For manuscript title:**

### Uneven declines between corals and cryptobenthic fish symbionts from multiple disturbances

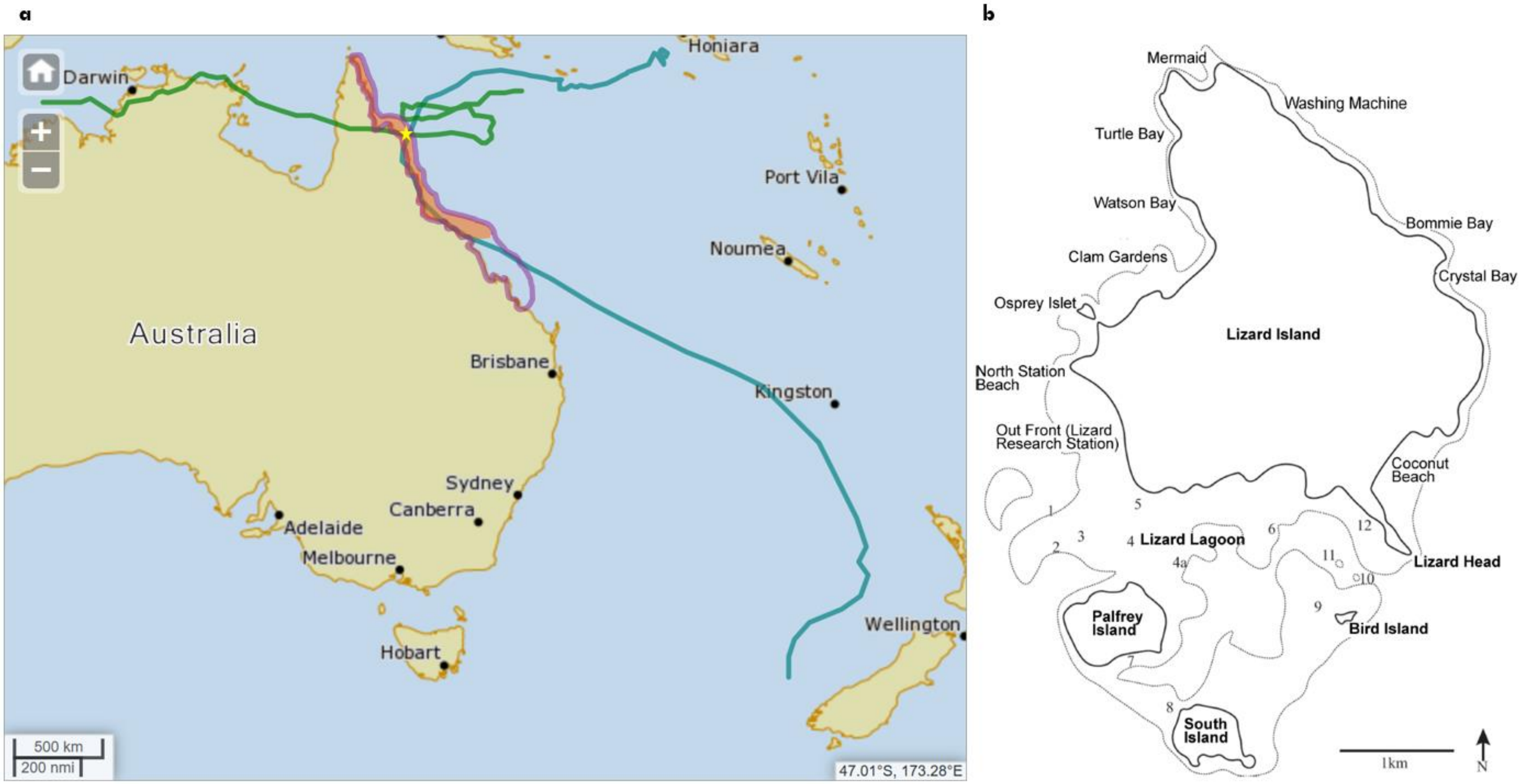


**Supplementary Figure 1. Map and location of consecutive climate disturbances affecting Lizard Island, QLD, Australia. a** Map of Australia with cyclone tracks of Ita 2014 (blue line) and Nathan 2015 (green line) from the Australian Government Bureau of Meteorology (http://www.bom.gov.au/cyclone/history/tracks/beta/?region=sh), with areas overlayed to show impacts of coral bleaching from 2016 (orange shaded area) and 2017 (purple outlined area). Lizard Island is highlighted with yellow star. Bleaching areas were adapted from two papers[15,16]. **b** Map of Lizard Island (-14.687264, 145.447039). Numbers represent discrete sites as follows: Big Vickey's Reef (1); Vickey's Reef (2); Horseshoe Reef (3); Palfrey Reef (4±4a); Loomis Reef (5); Trawler (6); Picnic Beach (7); Ghost Beach (8); Bird Island Reef (9); Entrance Bommie (10); Bird Bommie (11); Lizard Head (12).

**Supplementary Table 1. Statistical output of all univariate and multivariate analyses for changes in *Acropora* corals and coral-dwelling *Gobiodon* gobies following multiple disturbances.** Corals and gobies were surveyed before (Feb 2014) and after each cyclone (Jan-Feb 2015, Jan-Feb 2016) and consecutive bleaching (Feb-Mar 2018) around Lizard Island, Great Barrier Reef, Australia. CI means confidence interval.

| Response Variable | Model | Predictor variable | Factor Type | df | Model Statistic | Test-value | p-value | R-squared | Pairwise Comparison | Pairwise Statistic | Test-value | P-value | Lower 95% CI | Upper 95% CI |
|---|---|---|---|---|---|---|---|---|---|---|---|---|---|---|
| Coral Richness | GLMM poisson | Year | fixed | 3 | $\chi^2$ (chi-squared) | 35.418 | < 0.0001 | 0.147 marginal | 2014, 2015 | t-ratio | -0.34 | 0.9864 | -0.307 | 0.236 |
| | | Site | random | | | | | | 2014, 2016 | | 2.658 | 0.0412 | 0.007 | 0.53 |
| | | | | | | | | | 2014, 2018 | | 4.875 | 0.0001 | 0.349 | 1.137 |
| | | | | Sample | size | n = 279 | | | 2015, 2016 | | 3.203 | 0.0083 | 0.059 | 0.55 |
| | | | | Outliers | removed | n = 5 | | | 2015, 2018 | | 5.09 | <.0001 | 0.383 | 1.174 |
| | | | | | | | | | 2016, 2018 | | 3.18 | 0.0089 | 0.089 | 0.86 |
| Average Coral Diameter log-transformed | LMM | Year | fixed | 3 | F-value | 67.207 | < 0.0001 | 0.5874422 conditional | 2014, 2015 | t-ratio | 6.6 | <.0001 | 0.19419 | 0.445 |
| | | Site | random | | | | | | 2014, 2016 | | 9.572 | <.0001 | 0.31755 | 0.553 |
| | | | | | | | | | 2014, 2018 | | 13.334 | <.0001 | 0.72185 | 1.07 |
| | | | | Sample | size | n = 244 | | | 2015, 2016 | | 2.797 | 0.0285 | 0.00863 | 0.223 |
| | | | | Outliers | removed | n = 7 | | | 2015, 2018 | | 8.406 | <.0001 | 0.39888 | 0.754 |
| | | | | | | | | | 2016, 2018 | | 7.086 | <.0001 | 0.2923 | 0.629 |
| Coral Assemblage | PERMANOVA | Year | fixed | (3,259) | pseudo-F | 10.361 | 0.0001 | N/A | 2014, 2015 | t-value | 5.8509 | 0.0001 | N/A | N/A |
| | | Site | Random | (16,259) | | 2.7032 | 0.0001 | | 2014, 2016 | | 4.0838 | 0.0001 | | |
| | | | | | | | | | 2014, 2018 | | 4.6241 | 0.0001 | | |
| | | | | Sample | size | n = 279 | | | 2015, 2016 | | 1.7032 | 0.0092 | | |
| | | | | | | | | | 2015, 2018 | | 1.704 | 0.0066 | | |
| | | | | | | | | | 2016, 2018 | | 2.1477 | 0.0002 | | |
| Goby Richness | GLMM poisson | Year | fixed | 3 | $\chi^2$ (chi-squared) | 99.332 | < 0.0001 | 0.524 marginal | 2014, 2015 | t-ratio | 0.195 | 0.9974 | -0.249 | 0.289 |
| | | Site | random | | | | | | 2014, 2016 | | 2.545 | 0.0555 | -0.004 | 0.512 |
| | | | | | | | | | 2014, 2018 | | 7.999 | <.0001 | 1.277 | 2.496 |
| | | | | Sample | size | n = 279 | | | 2015, 2016 | | 2.512 | 0.0602 | -0.007 | 0.475 |
| | | | | Outliers | removed | n = 3 | | | 2015, 2018 | | 7.886 | <.0001 | 1.254 | 2.478 |
| | | | | | | | | | 2016, 2018 | | 6.975 | <.0001 | 1.027 | 2.237 |
| Average Goby Group Size log-transformed | LMM | Year | fixed | 3 | F-value | 12,877 | < 0.0001 | 0.2007437 conditional | 2014, 2015 | t-ratio | 4.308 | 0.0001 | 0.1420 | 0.569 |
| | | Site | random | | | | | | 2014, 2016 | | 4.645 | <.0001 | 0.1603 | 0.564 |
| | | | | | | | | | 2014, 2018 | | 5.088 | <.0001 | 0.3688 | 1.133 |
| | | | | Sample | size | n = 230 | | | 2015, 2016 | | 0.088 | 0.9998 | -0.1802 | 0.193 |
| | | | | Outliers | removed | n = 3 | | | 2015, 2018 | | 2.683 | 0.0389 | 0.0140 | 0.776 |
| | | | | | | | | | 2016, 2018 | | 2.709 | 0.0364 | 0.0172 | 0.760 |
| Percent Occupied Coral | LMM | Year | fixed | 3 | F-value | 79.009 | < 0.0001 | 0.6253415 conditional | 2014, 2015 | t-ratio | 5.876 | <.0001 | 10.88 | 28 |
| | | Site | random | | | | | | 2014, 2016 | | 4.394 | <.0001 | 5.55 | 21.5 |
| | | | | | | | | | 2014, 2018 | | 15.143 | <.0001 | 59.63 | 84.2 |
| | | | | Sample | size | n = 244 | | | 2015, 2016 | | -2.066 | 0.1674 | -13.37 | 1.5 |
| | | | | Outliers | removed | n = 6 | | | 2015, 2018 | | 10.739 | <.0001 | 39.84 | 65.1 |
| | | | | | | | | | 2016, 2018 | | 12.625 | <.0001 | 46.45 | 70.4 |
| Goby Assemblage | PERMANOVA | Year | fixed | (3,259) | pseudo-F | 5.8348 | 0.0001 | N/A | 2014, 2015 | t-value | 1.709 | 0.0061 | N/A | N/A |
| | | Site | Random | (16,259) | | 2.2043 | 0.0001 | | 2014, 2016 | | 1.777 | 0.0042 | | |
| | | | | | | | | | 2014, 2018 | | 2.9856 | 0.0001 | | |
| | | | | Sample | size | n = 279 | | | 2015, 2016 | | 2.1039 | 0.0008 | | |
| | | | | | | | | | 2015, 2018 | | 3.6387 | 0.0001 | | |
| | | | | | | | | | 2016, 2018 | | 2.7908 | 0.0001 | | |
